## Supplemental Figures and Datasets for "Characterisation of the RNA-interference pathway as a Tool for Genetics in the Nascent Phototrophic Endosymbiosis, *Paramecium bursaria*"

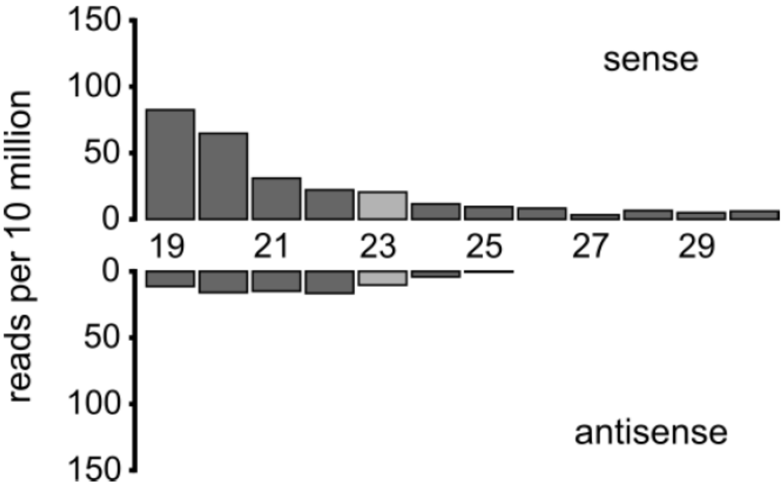


**Figure S1. Free-living Algal sRNA Abundance.** Size distribution of sRNA mapped to endosymbiont mRNA, extracted from a free-living algal culture (*Chlorella variabilis* NC64A, a known endosymbiont isolated from *P. bursaria*). Data are represented as mean ± SD of three biological replicates, and normalised to total reads per sample. Curated 'endosymbiont' transcript bins used for sRNA mapping are available at Figshare ([https://doi.org/10.6084/m9.figshare.12301736.v2](https://doi.org/10.6084/m9.figshare.12301736.v2)).

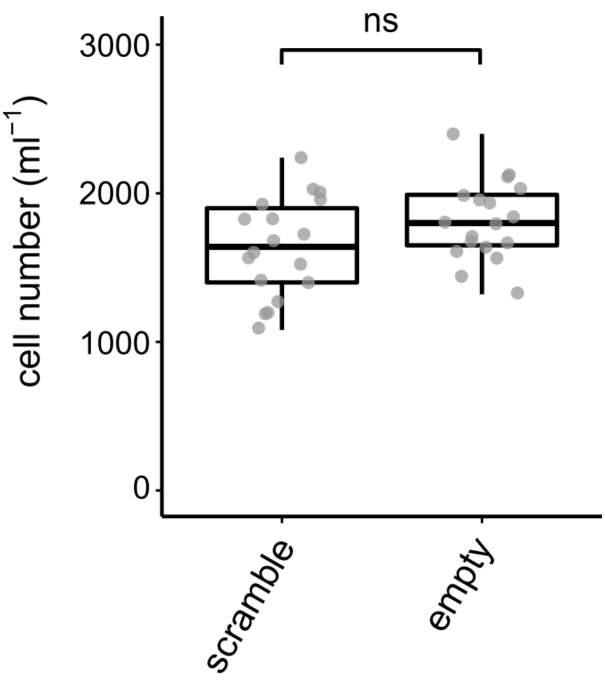

**Figure S2. Confirmation of the Null Effect of Scramble dsRNA Exposure.** *P. bursaria* cell number after feeding for 12 days with *E. coli* expressing non-hit 'scramble' dsRNA, or an empty vector control. Boxplot data are represented as max, upper quartile (Q3), mean, lower quartile (Q1) and min values of six biological replicates. Significance calculated using a generalized linear model with quasi-Poisson distribution.

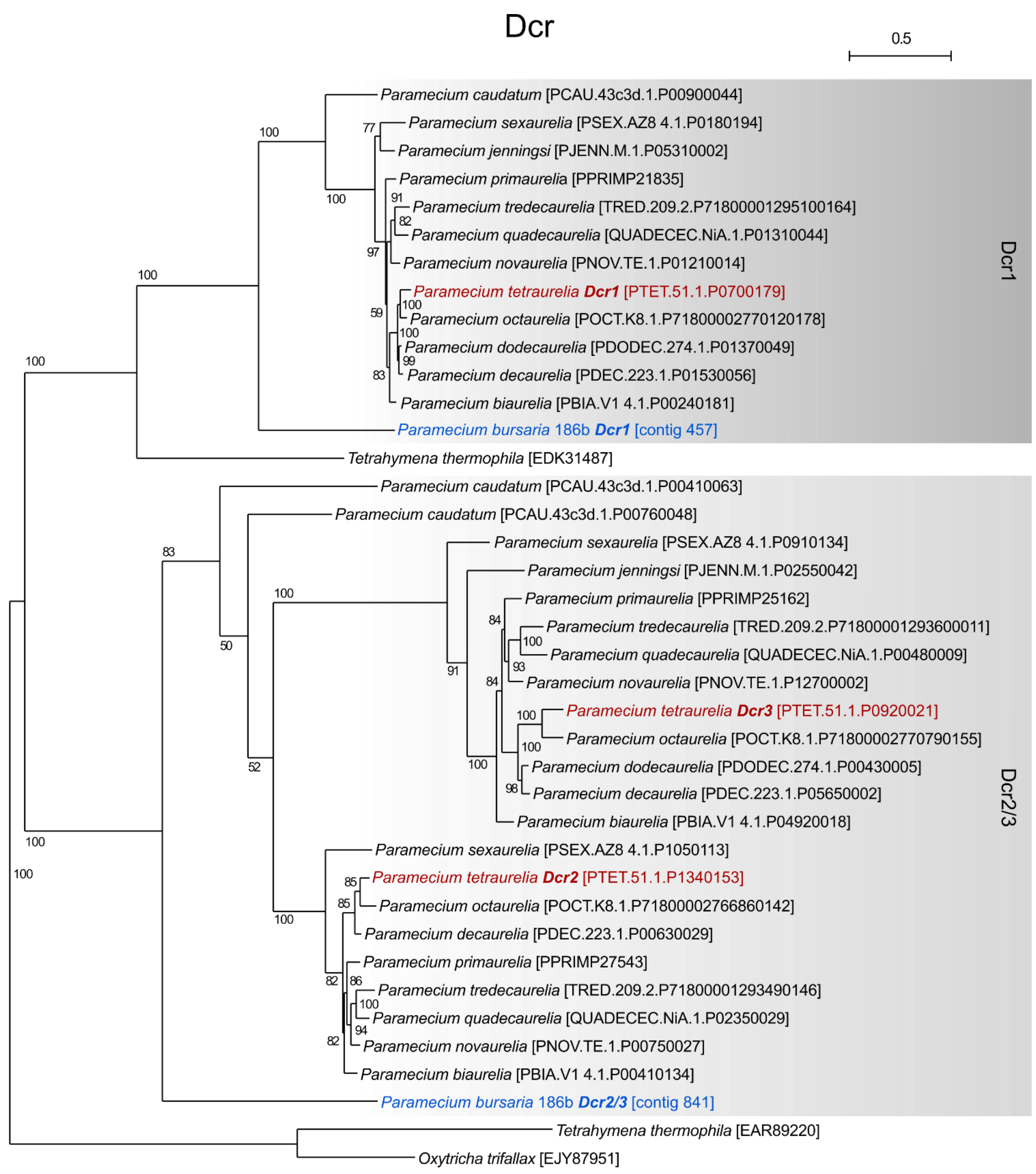
Dcr
0.5
Paramecium caudatum [PCAU.43c3d.1.P00900044]
Paramecium sexaurelia [PSEX.AZ8 4.1.P0180194]
Paramecium jenningsi [PJENN.M.1.P05310002]
Paramecium primaurelia [PPRIMP21835]
Paramecium tredecaurelia [TRED.209.2.P71800001295100164]
Paramecium quadecaurelia [QUADECEC.NiA.1.P01310044]
Paramecium novaurelia [PNOV.TE.1.P01210014]
Paramecium tetraurelia Dcr1 [PTET.51.1.P0700179]
Paramecium octaurelia [POCT.K8.1.P71800002770120178]
Paramecium dodecaurelia [PDODEC.274.1.P01370049]
Paramecium decaurelia [PDEC.223.1.P01530056]
Paramecium biaurelia [PBIA.V1 4.1.P00240181]
Paramecium bursaria 186b Dcr1 [contig 457]
Dcr1
Tetrahymena thermophila [EDK31487]
Paramecium caudatum [PCAU.43c3d.1.P00410063]
Paramecium caudatum [PCAU.43c3d.1.P00760048]
Paramecium sexaurelia [PSEX.AZ8 4.1.P0910134]
Paramecium jenningsi [PJENN.M.1.P02550042]
Paramecium primaurelia [PPRIMP25162]
Paramecium tredecaurelia [TRED.209.2.P71800001293600011]
Paramecium quadecaurelia [QUADECEC.NiA.1.P00480009]
Paramecium novaurelia [PNOV.TE.1.P12700002]
Paramecium tetraurelia Dcr3 [PTET.51.1.P0920021]
Paramecium octaurelia [POCT.K8.1.P71800002770790155]
Paramecium dodecaurelia [PDODEC.274.1.P00430005]
Paramecium decaurelia [PDEC.223.1.P05650002]
Paramecium biaurelia [PBIA.V1 4.1.P04920018]
Paramecium sexaurelia [PSEX.AZ8 4.1.P1050113]
Paramecium tetraurelia Dcr2 [PTET.51.1.P1340153]
Paramecium octaurelia [POCT.K8.1.P71800002766860142]
Paramecium decaurelia [PDEC.223.1.P00630029]
Paramecium primaurelia [PPRIMP27543]
Paramecium tredecaurelia [TRED.209.2.P71800001293490146]
Paramecium quadecaurelia [QUADECEC.NiA.1.P02350029]
Paramecium novaurelia [PNOV.TE.1.P00750027]
Paramecium biaurelia [PBIA.V1 4.1.P00410134]
Paramecium bursaria 186b Dcr2/3 [contig 841]
Dcr2/3
Tetrahymena thermophila [EAR89220]
Oxytricha trifallax [EJY87951]

**Dataset S1.** *Dcr* phylogeny (based on 685 sampled aligned amino acid sites) calculated using iqtree with a JTT+F+G4 best fit substitution model chosen according to Bayesian Inference Criterion (BIC), and with 1,000 non-parametric bootstrap replicates. Classification of *P. bursaria* Dicer sequences demonstrates two clear orthologues: a single *Dcr1* orthologue, and a single un-duplicated *Dcr2/3* orthologue. Scale bars for all phylogenies indicate average substitutions per site. Red indicates annotated *P. tetraurelia* query seed sequences[1] used to search genome/transcriptome databases (02/12/2020), blue indicates sequences identified from the respective *P. bursaria* datasets. Genomic sequence data for *P. bursaria* 186b was used to supplement the partial transcriptome data available. Nucleotide sequence (https://doi.org/10.6084/m9.figshare.13387811.v1) and amino acid alignment data (https://doi.org/10.6084/m9.figshare.13387631.v1) for putative *P. bursaria* homologues are available at Figshare.

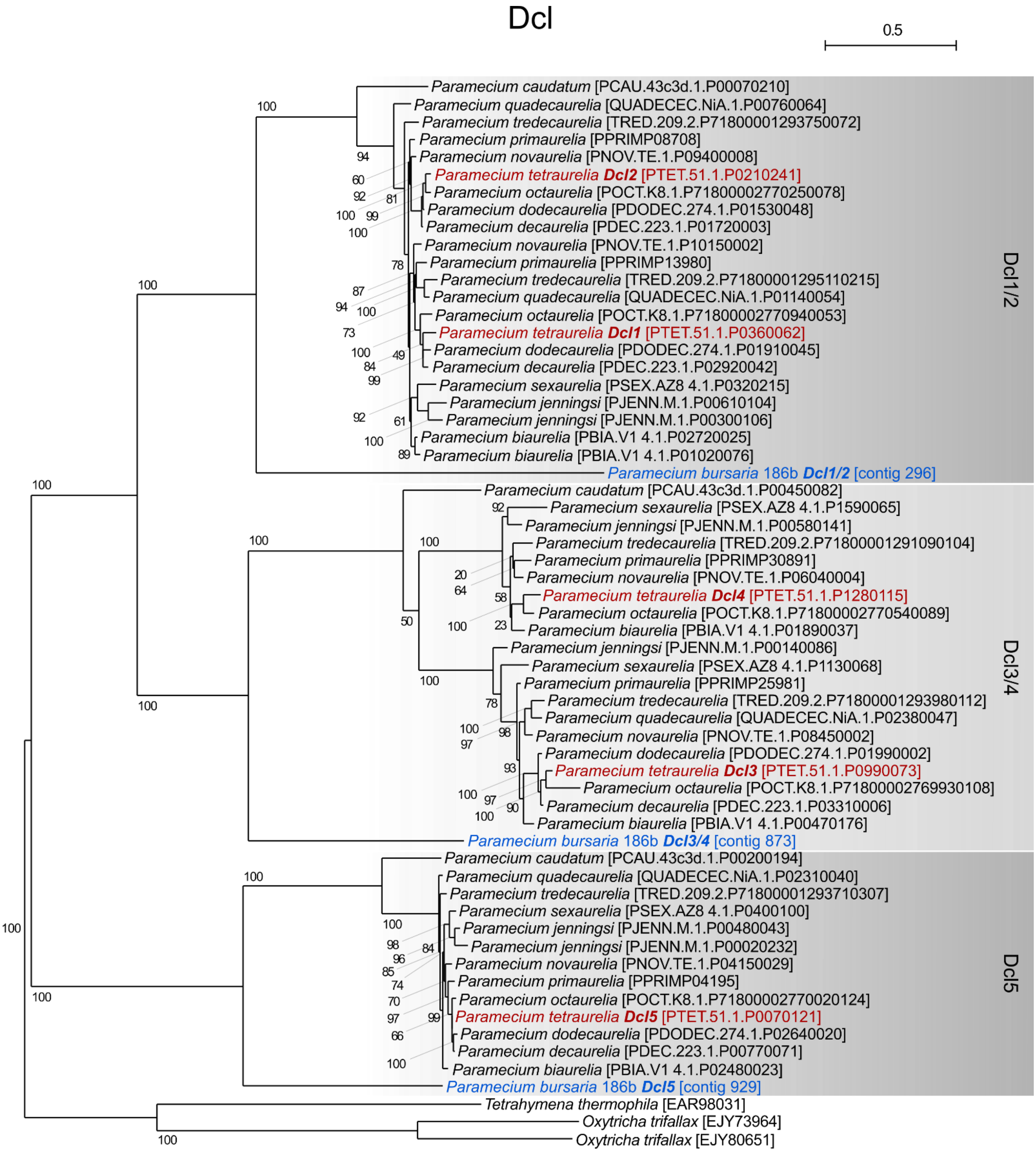


**Dataset S2.** *Dcl* phylogeny (based on 320 sampled aligned amino acid sites) calculated using iqtree with a LG+F+I+G4 best fit substitution model chosen according to Bayesian Inference Criterion (BIC), and with 1,000 non-parametric bootstrap replicates. Classification of *P. bursaria* Dicer-like sequences demonstrates three clear orthologues: a single *Dcl5* orthologue, and two un-duplicated *Dcl1/2* and *Dcl3/4* orthologues. Genomic sequence data for *P. bursaria* 186b was used to supplement the partial transcriptome data available. Nucleotide sequence (https://doi.org/10.6084/m9.figshare.13387811.v1) and amino acid alignment data (https://doi.org/10.6084/m9.figshare.13387631.v1) for putative *P. bursaria* homologues are available at Figshare.

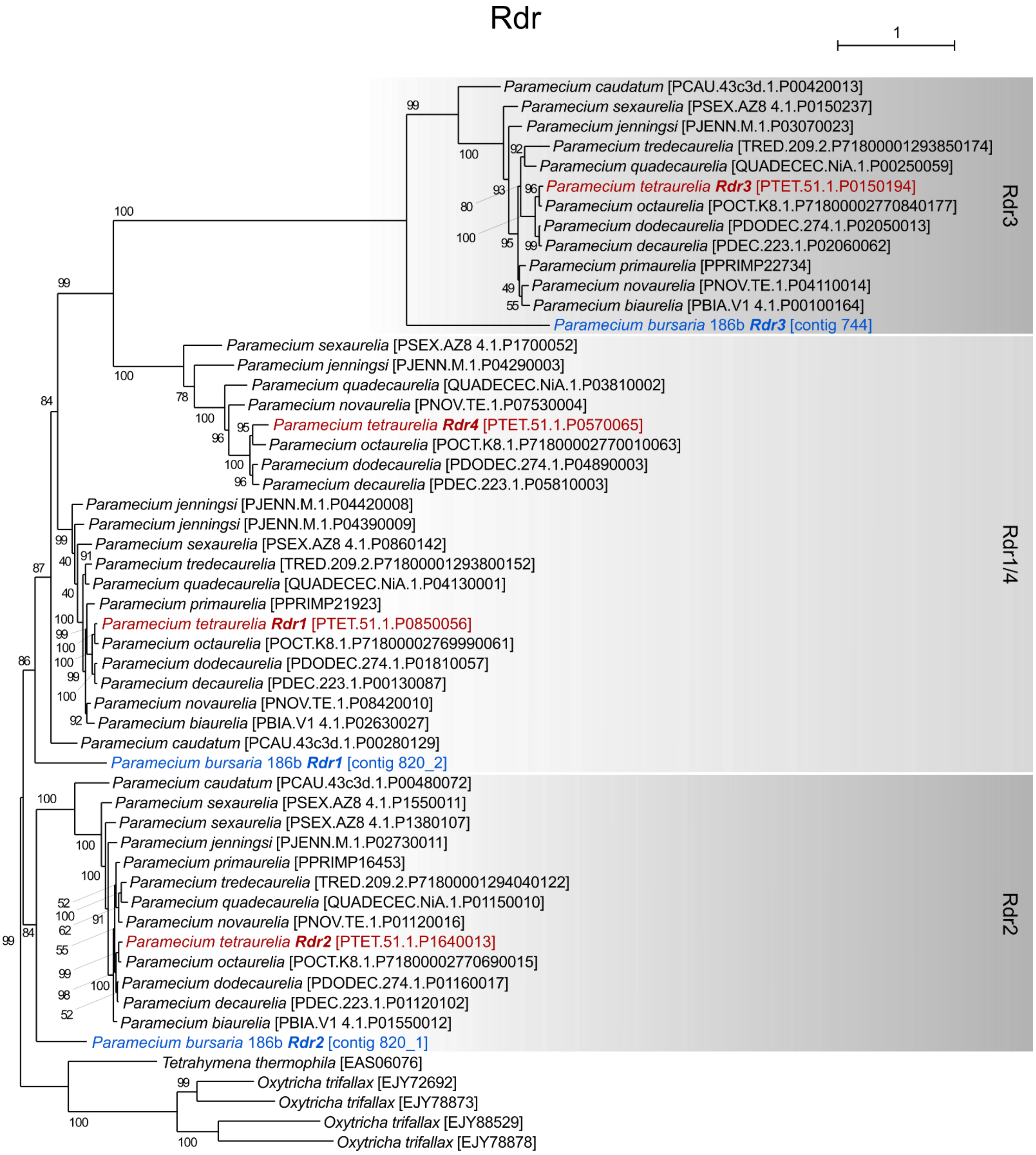
Rdr
1
Paramecium caudatum [PCAU.43c3d.1.P00420013]
Paramecium sexaurelia [PSEX.AZ8 4.1.P0150237]
Paramecium jenningsi [PJENN.M.1.P03070023]
Paramecium tredecaurelia [TRED.209.2.P71800001293850174]
Paramecium quadecaurelia [QUADECEC.NiA.1.P00250059]
Paramecium tetraurelia Rdr3 [PTET.51.1.P0150194]
Paramecium octaurelia [POCT.K8.1.P71800002770840177]
Paramecium dodecaurelia [PDODEC.274.1.P02050013]
Paramecium decaurelia [PDEC.223.1.P02060062]
Paramecium primaurelia [PPRIMP22734]
Paramecium novaurelia [PNOV.TE.1.P04110014]
Paramecium biaurelia [PBIA.V1 4.1.P00100164]
Paramecium bursaria 186b Rdr3 [contig 744]
Rdr3
Paramecium sexaurelia [PSEX.AZ8 4.1.P1700052]
Paramecium jenningsi [PJENN.M.1.P04290003]
Paramecium quadecaurelia [QUADECEC.NiA.1.P03810002]
Paramecium novaurelia [PNOV.TE.1.P07530004]
Paramecium tetraurelia Rdr4 [PTET.51.1.P0570065]
Paramecium octaurelia [POCT.K8.1.P71800002770010063]
Paramecium dodecaurelia [PDODEC.274.1.P04890003]
Paramecium decaurelia [PDEC.223.1.P05810003]
Paramecium jenningsi [PJENN.M.1.P04420008]
Paramecium jenningsi [PJENN.M.1.P04390009]
Paramecium sexaurelia [PSEX.AZ8 4.1.P0860142]
Paramecium tredecaurelia [TRED.209.2.P71800001293800152]
Paramecium quadecaurelia [QUADECEC.NiA.1.P04130001]
Paramecium primaurelia [PPRIMP21923]
Paramecium tetraurelia Rdr1 [PTET.51.1.P0850056]
Paramecium octaurelia [POCT.K8.1.P71800002769990061]
Paramecium dodecaurelia [PDODEC.274.1.P01810057]
Paramecium decaurelia [PDEC.223.1.P00130087]
Paramecium novaurelia [PNOV.TE.1.P08420010]
Paramecium biaurelia [PBIA.V1 4.1.P02630027]
Paramecium caudatum [PCAU.43c3d.1.P00280129]
Paramecium bursaria 186b Rdr1 [contig 820_2]
Rdr1/4
Paramecium caudatum [PCAU.43c3d.1.P00480072]
Paramecium sexaurelia [PSEX.AZ8 4.1.P1550011]
Paramecium sexaurelia [PSEX.AZ8 4.1.P1380107]
Paramecium jenningsi [PJENN.M.1.P02730011]
Paramecium primaurelia [PPRIMP16453]
Paramecium tredecaurelia [TRED.209.2.P71800001294040122]
Paramecium quadecaurelia [QUADECEC.NiA.1.P01150010]
Paramecium novaurelia [PNOV.TE.1.P01120016]
Paramecium tetraurelia Rdr2 [PTET.51.1.P1640013]
Paramecium octaurelia [POCT.K8.1.P71800002770690015]
Paramecium dodecaurelia [PDODEC.274.1.P01160017]
Paramecium decaurelia [PDEC.223.1.P01120102]
Paramecium biaurelia [PBIA.V1 4.1.P01550012]
Paramecium bursaria 186b Rdr2 [contig 820_1]
Rdr2
Tetrahymena thermophila [EAS06076]
Oxytricha trifallax [EJY72692]
Oxytricha trifallax [EJY78873]
Oxytricha trifallax [EJY88529]
Oxytricha trifallax [EJY78878]

**Dataset S3.** *Rdr* phylogeny (based on 406 sampled aligned amino acid sites) calculated using iqtree with a LG+F+I+G4 best fit substitution model chosen according to Bayesian Inference Criterion (BIC), and with 1,000 non-parametric bootstrap replicates. Classification of *P. bursaria Rdr* sequences relative to *Rdr4* is ambiguous, but demonstrates three clear orthologues: a single *Rdr1* orthologue, a single *Rdr3* orthologue, and a putative un-duplicated *Rdr1/4* orthologue. Genomic sequence data for *P. bursaria* 186b was used to supplement the partial transcriptome data available. Nucleotide sequence (https://doi.org/10.6084/m9.figshare.13387811.v1) and amino acid alignment data (https://doi.org/10.6084/m9.figshare.13387631.v1) for putative *P. bursaria* homologues are available at Figshare.

a

PiwiA

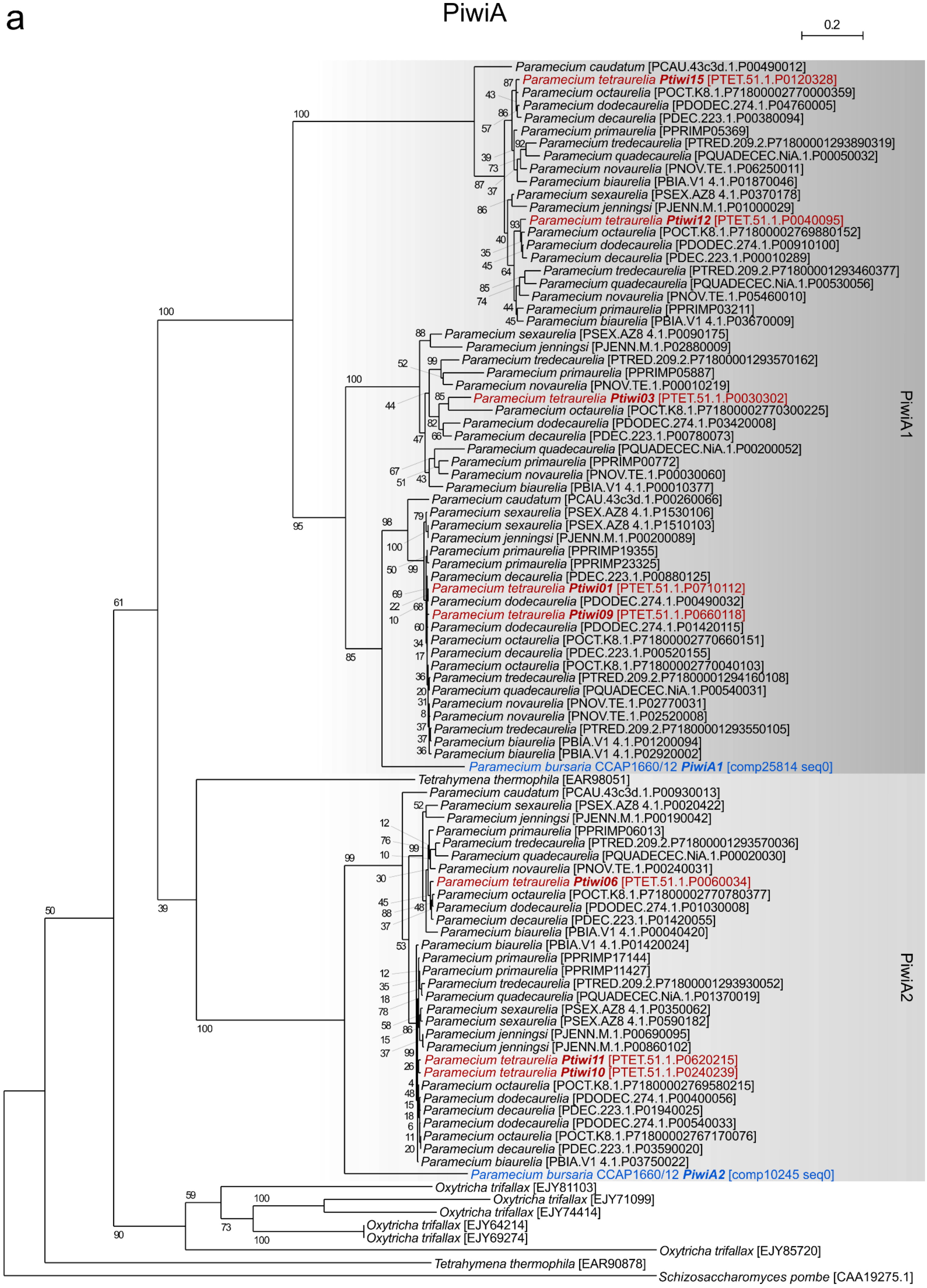

0.2
Paramecium caudatum [PCAU.43c3d.1.P00490012]
Paramecium tetraurelia Ptiwi15 [PTET.51.1.P0120328]
Paramecium octaurelia [POCT.K8.1.P71800002770000359]
Paramecium dodecaurelia [PDODEC.274.1.P04760005]
Paramecium decaurelia [PDEC.223.1.P00380094]
Paramecium primaurelia [PPRIMP05369]
Paramecium tredecaurelia [PTRED.209.2.P71800001293890319]
Paramecium quadecaurelia [PQUADECEC.NiA.1.P00050032]
Paramecium novaurelia [PNOV.TE.1.P06250011]
Paramecium biaurelia [PBIA.V1 4.1.P01870046]
Paramecium sexaurelia [PSEX.AZ8 4.1.P0370178]
Paramecium jenningsi [PJENN.M.1.P01000029]
Paramecium tetraurelia Ptiwi12 [PTET.51.1.P0040095]
Paramecium octaurelia [POCT.K8.1.P71800002769880152]
Paramecium dodecaurelia [PDODEC.274.1.P00910100]
Paramecium decaurelia [PDEC.223.1.P00010289]
Paramecium tredecaurelia [PTRED.209.2.P71800001293460377]
Paramecium quadecaurelia [PQUADECEC.NiA.1.P00530056]
Paramecium novaurelia [PNOV.TE.1.P05460010]
Paramecium primaurelia [PPRIMP03211]
Paramecium biaurelia [PBIA.V1 4.1.P03670009]
Paramecium sexaurelia [PSEX.AZ8 4.1.P0090175]
Paramecium jenningsi [PJENN.M.1.P02880009]
Paramecium tredecaurelia [PTRED.209.2.P71800001293570162]
Paramecium primaurelia [PPRIMP05887]
Paramecium novaurelia [PNOV.TE.1.P00010219]
Paramecium tetraurelia Ptiwi03 [PTET.51.1.P0030302]
Paramecium octaurelia [POCT.K8.1.P71800002770300225]
Paramecium dodecaurelia [PDODEC.274.1.P03420008]
Paramecium decaurelia [PDEC.223.1.P00780073]
Paramecium quadecaurelia [PQUADECEC.NiA.1.P00200052]
Paramecium primaurelia [PPRIMP00772]
Paramecium novaurelia [PNOV.TE.1.P00030060]
Paramecium biaurelia [PBIA.V1 4.1.P00010377]
Paramecium caudatum [PCAU.43c3d.1.P00260066]
Paramecium sexaurelia [PSEX.AZ8 4.1.P1530106]
Paramecium sexaurelia [PSEX.AZ8 4.1.P1510103]
Paramecium jenningsi [PJENN.M.1.P00200089]
Paramecium primaurelia [PPRIMP19355]
Paramecium primaurelia [PPRIMP23325]
Paramecium decaurelia [PDEC.223.1.P00880125]
Paramecium tetraurelia Ptiwi01 [PTET.51.1.P0710112]
Paramecium dodecaurelia [PDODEC.274.1.P00490032]
Paramecium tetraurelia Ptiwi09 [PTET.51.1.P0660118]
Paramecium dodecaurelia [PDODEC.274.1.P01420115]
Paramecium octaurelia [POCT.K8.1.P71800002770660151]
Paramecium decaurelia [PDEC.223.1.P00520155]
Paramecium octaurelia [POCT.K8.1.P71800002770040103]
Paramecium tredecaurelia [PTRED.209.2.P71800001294160108]
Paramecium quadecaurelia [PQUADECEC.NiA.1.P00540031]
Paramecium novaurelia [PNOV.TE.1.P02770031]
Paramecium novaurelia [PNOV.TE.1.P02520008]
Paramecium tredecaurelia [PTRED.209.2.P71800001293550105]
Paramecium biaurelia [PBIA.V1 4.1.P01200094]
Paramecium biaurelia [PBIA.V1 4.1.P02920002]
Paramecium bursaria CCAP1660/12 PiwiA1 [comp25814 seq0]
PiwiA1
Tetrahymena thermophila [EAR98051]
Paramecium caudatum [PCAU.43c3d.1.P00930013]
Paramecium sexaurelia [PSEX.AZ8 4.1.P0020422]
Paramecium jenningsi [PJENN.M.1.P00190042]
Paramecium primaurelia [PPRIMP06013]
Paramecium tredecaurelia [PTRED.209.2.P71800001293570036]
Paramecium quadecaurelia [PQUADECEC.NiA.1.P00020030]
Paramecium novaurelia [PNOV.TE.1.P00240031]
Paramecium tetraurelia Ptiwi06 [PTET.51.1.P0060034]
Paramecium octaurelia [POCT.K8.1.P71800002770780377]
Paramecium dodecaurelia [PDODEC.274.1.P01030008]
Paramecium decaurelia [PDEC.223.1.P01420055]
Paramecium biaurelia [PBIA.V1 4.1.P00040420]
Paramecium biaurelia [PBIA.V1 4.1.P01420024]
Paramecium primaurelia [PPRIMP17144]
Paramecium primaurelia [PPRIMP11427]
Paramecium tredecaurelia [PTRED.209.2.P71800001293930052]
Paramecium quadecaurelia [PQUADECEC.NiA.1.P01370019]
Paramecium sexaurelia [PSEX.AZ8 4.1.P0350062]
Paramecium sexaurelia [PSEX.AZ8 4.1.P0590182]
Paramecium jenningsi [PJENN.M.1.P00690095]
Paramecium jenningsi [PJENN.M.1.P00860102]
Paramecium tetraurelia Ptiwi11 [PTET.51.1.P0620215]
Paramecium tetraurelia Ptiwi10 [PTET.51.1.P0240239]
Paramecium octaurelia [POCT.K8.1.P71800002769580215]
Paramecium dodecaurelia [PDODEC.274.1.P00400056]
Paramecium decaurelia [PDEC.223.1.P01940025]
Paramecium dodecaurelia [PDODEC.274.1.P00540033]
Paramecium octaurelia [POCT.K8.1.P71800002767170076]
Paramecium decaurelia [PDEC.223.1.P03590020]
Paramecium biaurelia [PBIA.V1 4.1.P03750022]
Paramecium bursaria CCAP1660/12 PiwiA2 [comp10245 seq0]
PiwiA2
Oxytricha trifallax [EJY81103]
Oxytricha trifallax [EJY71099]
Oxytricha trifallax [EJY74414]
Oxytricha trifallax [EJY64214]
Oxytricha trifallax [EJY69274]
Oxytricha trifallax [EJY85720]
Tetrahymena thermophila [EAR90878]
Schizosaccharomyces pombe [CAA19275.1]

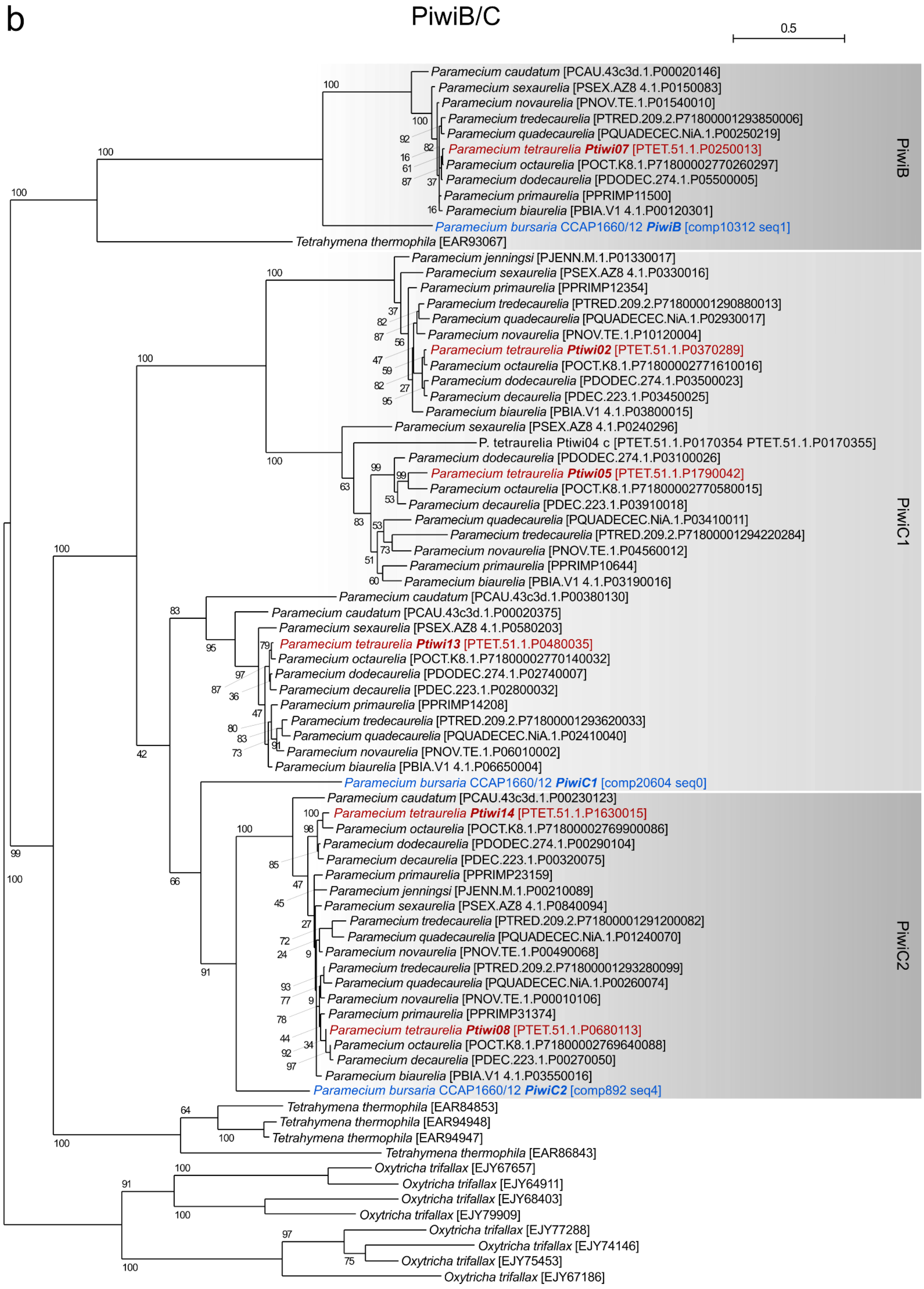
b
PiwiB/C
0.5
Paramecium caudatum [PCAU.43c3d.1.P00020146]
Paramecium sexaurelia [PSEX.AZ8 4.1.P0150083]
Paramecium novaurelia [PNOV.TE.1.P01540010]
Paramecium tredecaurelia [PTRED.209.2.P71800001293850006]
Paramecium quadecaurelia [PQUADECEC.NiA.1.P00250219]
Paramecium tetraurelia Ptiwi07 [PTET.51.1.P0250013]
Paramecium octaurelia [POCT.K8.1.P71800002770260297]
Paramecium dodecaurelia [PDODEC.274.1.P05500005]
Paramecium primaurelia [PPRIMP11500]
Paramecium biaurelia [PBIA.V1 4.1.P00120301]
Paramecium bursaria CCAP1660/12 PiwiB [comp10312 seq1]
Tetrahymena thermophila [EAR93067]
PiwiB
Paramecium jenningsi [PJENN.M.1.P01330017]
Paramecium sexaurelia [PSEX.AZ8 4.1.P0330016]
Paramecium primaurelia [PPRIMP12354]
Paramecium tredecaurelia [PTRED.209.2.P71800001290880013]
Paramecium quadecaurelia [PQUADECEC.NiA.1.P02930017]
Paramecium novaurelia [PNOV.TE.1.P10120004]
Paramecium tetraurelia Ptiwi02 [PTET.51.1.P0370289]
Paramecium octaurelia [POCT.K8.1.P71800002771610016]
Paramecium dodecaurelia [PDODEC.274.1.P03500023]
Paramecium decaurelia [PDEC.223.1.P03450025]
Paramecium biaurelia [PBIA.V1 4.1.P03800015]
Paramecium sexaurelia [PSEX.AZ8 4.1.P0240296]
P. tetraurelia Ptiwi04 c [PTET.51.1.P0170354 PTET.51.1.P0170355]
Paramecium dodecaurelia [PDODEC.274.1.P03100026]
Paramecium tetraurelia Ptiwi05 [PTET.51.1.P1790042]
Paramecium octaurelia [POCT.K8.1.P71800002770580015]
Paramecium decaurelia [PDEC.223.1.P03910018]
Paramecium quadecaurelia [PQUADECEC.NiA.1.P03410011]
Paramecium tredecaurelia [PTRED.209.2.P71800001294220284]
Paramecium novaurelia [PNOV.TE.1.P04560012]
Paramecium primaurelia [PPRIMP10644]
Paramecium biaurelia [PBIA.V1 4.1.P03190016]
Paramecium caudatum [PCAU.43c3d.1.P00380130]
Paramecium caudatum [PCAU.43c3d.1.P00020375]
Paramecium sexaurelia [PSEX.AZ8 4.1.P0580203]
Paramecium tetraurelia Ptiwi13 [PTET.51.1.P0480035]
Paramecium octaurelia [POCT.K8.1.P71800002770140032]
Paramecium dodecaurelia [PDODEC.274.1.P02740007]
Paramecium decaurelia [PDEC.223.1.P02800032]
Paramecium primaurelia [PPRIMP14208]
Paramecium tredecaurelia [PTRED.209.2.P71800001293620033]
Paramecium quadecaurelia [PQUADECEC.NiA.1.P02410040]
Paramecium novaurelia [PNOV.TE.1.P06010002]
Paramecium biaurelia [PBIA.V1 4.1.P06650004]
Paramecium bursaria CCAP1660/12 PiwiC1 [comp20604 seq0]
PiwiC1
Paramecium caudatum [PCAU.43c3d.1.P00230123]
Paramecium tetraurelia Ptiwi14 [PTET.51.1.P1630015]
Paramecium octaurelia [POCT.K8.1.P71800002769900086]
Paramecium dodecaurelia [PDODEC.274.1.P00290104]
Paramecium decaurelia [PDEC.223.1.P00320075]
Paramecium primaurelia [PPRIMP23159]
Paramecium jenningsi [PJENN.M.1.P00210089]
Paramecium sexaurelia [PSEX.AZ8 4.1.P0840094]
Paramecium tredecaurelia [PTRED.209.2.P71800001291200082]
Paramecium quadecaurelia [PQUADECEC.NiA.1.P01240070]
Paramecium novaurelia [PNOV.TE.1.P00490068]
Paramecium tredecaurelia [PTRED.209.2.P71800001293280099]
Paramecium quadecaurelia [PQUADECEC.NiA.1.P00260074]
Paramecium novaurelia [PNOV.TE.1.P00010106]
Paramecium primaurelia [PPRIMP31374]
Paramecium tetraurelia Ptiwi08 [PTET.51.1.P0680113]
Paramecium octaurelia [POCT.K8.1.P71800002769640088]
Paramecium decaurelia [PDEC.223.1.P00270050]
Paramecium biaurelia [PBIA.V1 4.1.P03550016]
Paramecium bursaria CCAP1660/12 PiwiC2 [comp892 seq4]
PiwiC2
Tetrahymena thermophila [EAR84853]
Tetrahymena thermophila [EAR94948]
Tetrahymena thermophila [EAR94947]
Tetrahymena thermophila [EAR86843]
Oxytricha trifallax [EJY67657]
Oxytricha trifallax [EJY64911]
Oxytricha trifallax [EJY68403]
Oxytricha trifallax [EJY79909]
Oxytricha trifallax [EJY77288]
Oxytricha trifallax [EJY74146]
Oxytricha trifallax [EJY75453]
Oxytricha trifallax [EJY67186]

c

PiwiD

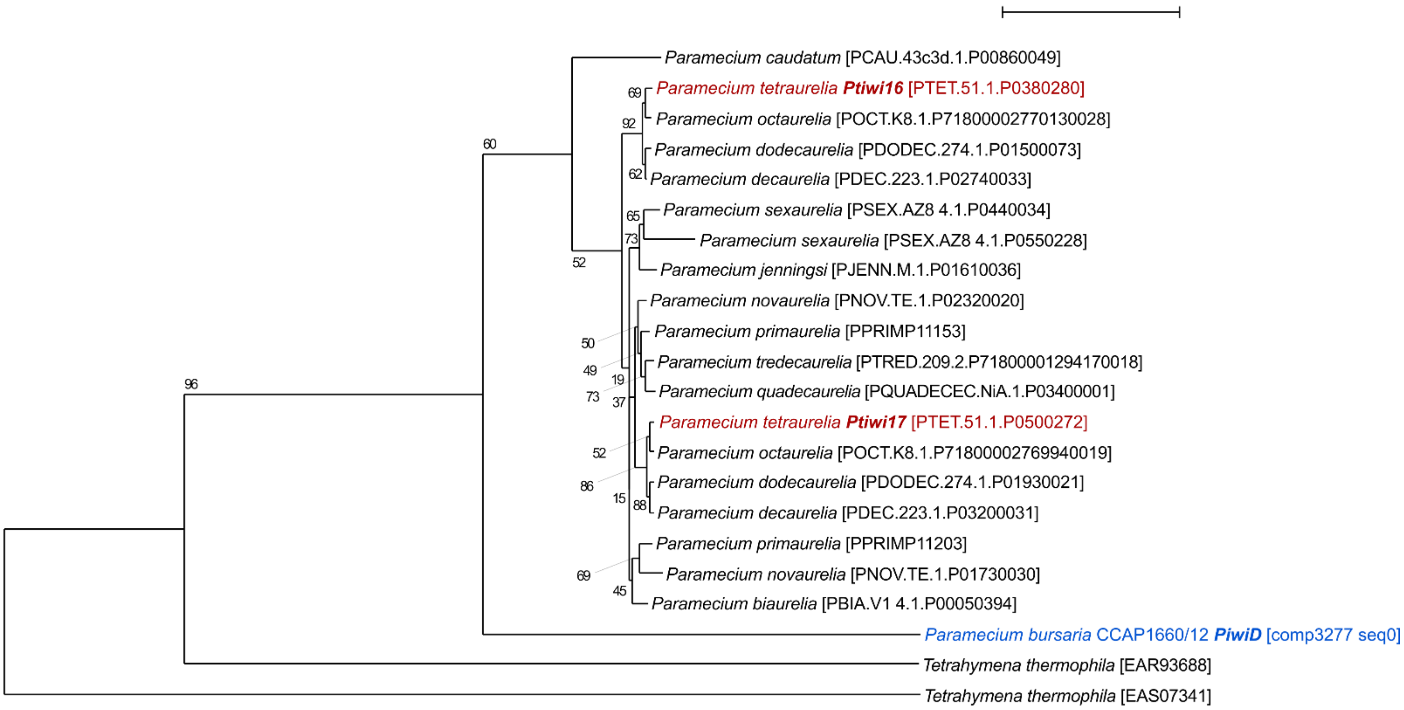


**Dataset S4.** (**a**) *PiwiA* phylogeny (based on 516 sampled aligned amino acid sites) calculated using iqtree with a LG+I+G best fit substitution model chosen according to Bayesian Inference Criterion (BIC), and with 1,000 non-parametric bootstrap replicates. (**b**) *PiwiB/C* phylogeny (based on 492 sampled aligned amino acid sites) calculated using iqtree with a LG+F+I+G4 best fit substitution model chosen according to Bayesian Inference Criterion (BIC), and with 1,000 non-parametric bootstrap replicates. (**c**) *PiwiD* phylogeny (based on 468 sampled aligned amino acid sites) calculated using iqtree with a JTTDCMut+F+R3 best fit substitution model chosen according to Bayesian Inference Criterion (BIC), and with 1,000 non-parametric bootstrap replicates. Classification of *P. bursaria* sequences relative to *P. tetraurelia Ptiwi* components is ambiguous, but demonstrates six clear orthologues: two putative unduplicated *PiwiA* (*Ptiwi01/03/09/12/15/06/10/11*) orthologues, a single *PiwiB* (*Ptiwi07*) orthologue, two putative unduplicated *PiwiC* (*Ptiwi02/04/05/13/08/14*) orthologues, and a single *PiwiD* (*Ptiwi16/17*) orthologue. Nucleotide sequence (https://doi.org/10.6084/m9.figshare.13387811.v1) and amino acid alignment data (https://doi.org/10.6084/m9.figshare.13387631.v1) for putative *P. bursaria* homologues are available at Figshare.

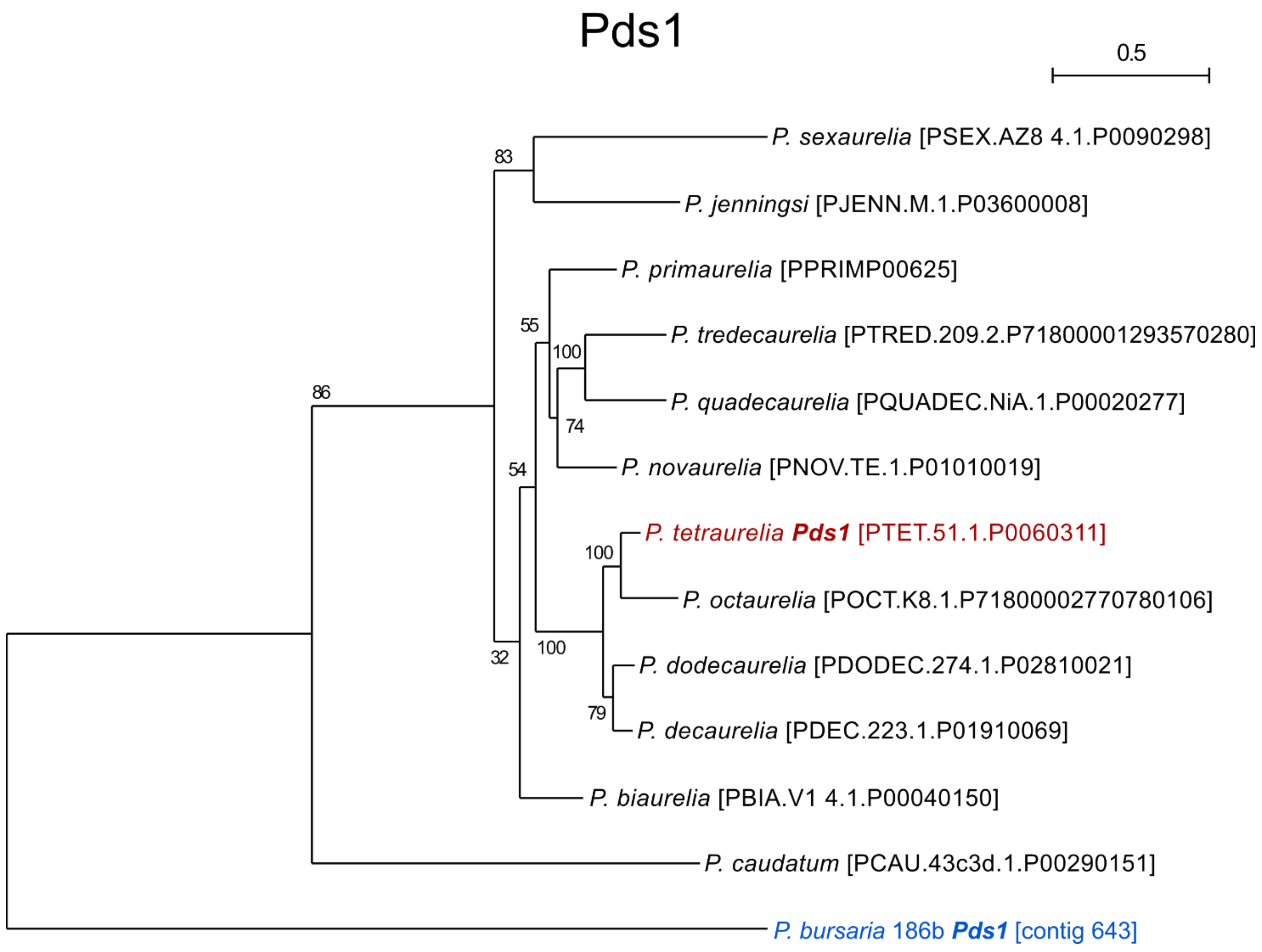


**Dataset S5.** *Pds1* phylogeny (based on 546 sampled aligned amino acid sites) calculated using iqtree with a JTT+F+G4 best fit substitution model chosen according to Bayesian Inference Criterion (BIC), and with 1,000 non-parametric bootstrap replicates. No homologue for *Pds1* could be identified in *T. thermophilia, O. trifallax,* or any broader ciliate dataset, indicating that this gene may be *Paramecium* specific. Nucleotide sequence (https://doi.org/10.6084/m9.figshare.13387811.v1) and amino acid alignment data (https://doi.org/10.6084/m9.figshare.13387631.v1) for putative *P. bursaria* homologues are available at Figshare.

### Cid

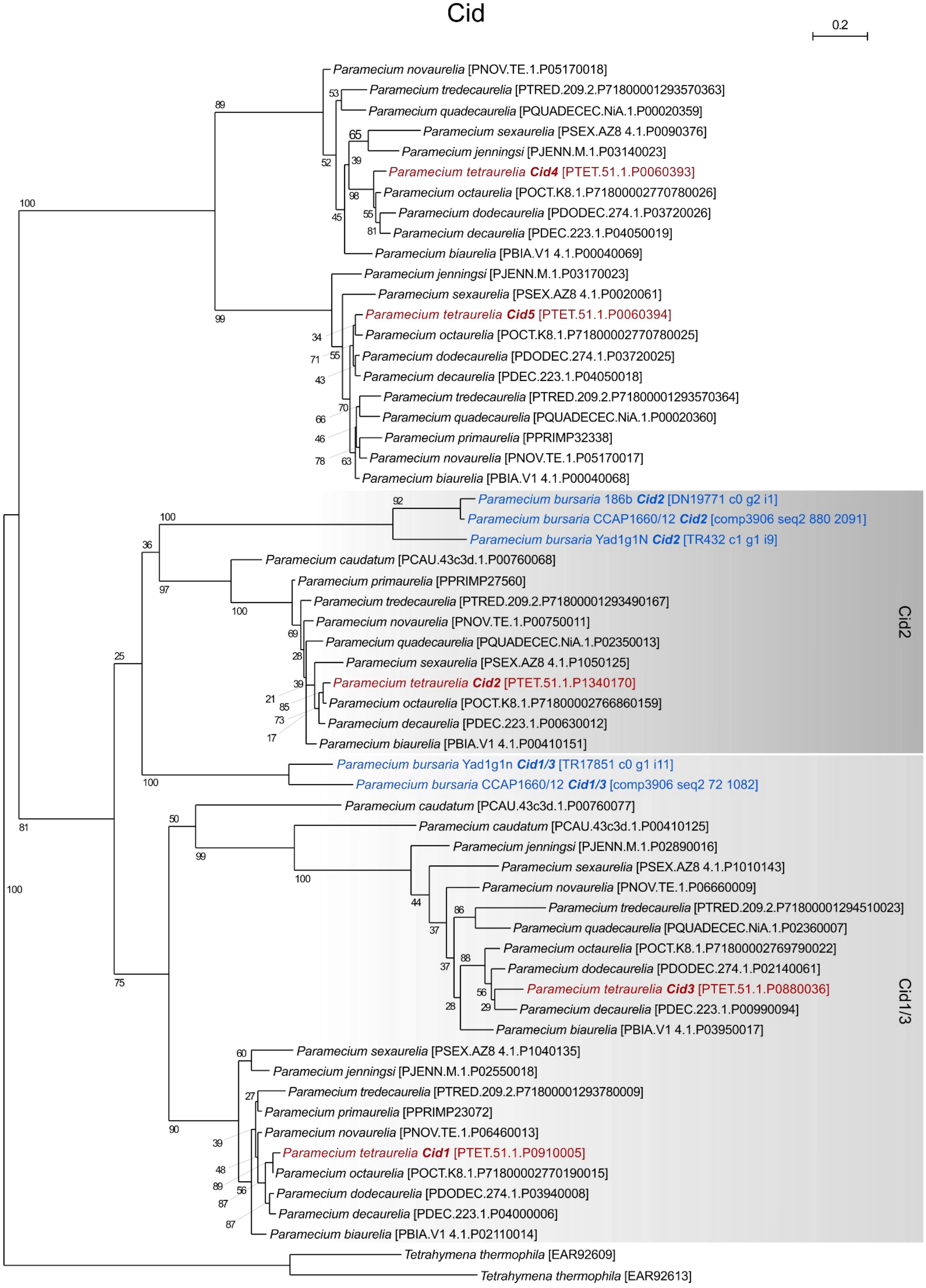

0.2
Paramecium novaurelia [PNOV.TE.1.P05170018]
Paramecium tredecaurelia [PTRED.209.2.P71800001293570363]
Paramecium quadecaurelia [PQUADECEC.NiA.1.P00020359]
Paramecium sexaurelia [PSEX.AZ8 4.1.P0090376]
Paramecium jenningsi [PJENN.M.1.P03140023]
Paramecium tetraurelia Cid4 [PTET.51.1.P0060393]
Paramecium octaurelia [POCT.K8.1.P71800002770780026]
Paramecium dodecaurelia [PDODEC.274.1.P03720026]
Paramecium decaurelia [PDEC.223.1.P04050019]
Paramecium biaurelia [PBIA.V1 4.1.P00040069]
Paramecium jenningsi [PJENN.M.1.P03170023]
Paramecium sexaurelia [PSEX.AZ8 4.1.P0020061]
Paramecium tetraurelia Cid5 [PTET.51.1.P0060394]
Paramecium octaurelia [POCT.K8.1.P71800002770780025]
Paramecium dodecaurelia [PDODEC.274.1.P03720025]
Paramecium decaurelia [PDEC.223.1.P04050018]
Paramecium tredecaurelia [PTRED.209.2.P71800001293570364]
Paramecium quadecaurelia [PQUADECEC.NiA.1.P00020360]
Paramecium primaurelia [PPRIMP32338]
Paramecium novaurelia [PNOV.TE.1.P05170017]
Paramecium biaurelia [PBIA.V1 4.1.P00040068]
Paramecium bursaria 186b Cid2 [DN19771 c0 g2 i1]
Paramecium bursaria CCAP1660/12 Cid2 [comp3906 seq2 880 2091]
Paramecium bursaria Yad1g1N Cid2 [TR432 c1 g1 i9]
Paramecium caudatum [PCAU.43c3d.1.P00760068]
Paramecium primaurelia [PPRIMP27560]
Paramecium tredecaurelia [PTRED.209.2.P71800001293490167]
Paramecium novaurelia [PNOV.TE.1.P00750011]
Paramecium quadecaurelia [PQUADECEC.NiA.1.P02350013]
Paramecium sexaurelia [PSEX.AZ8 4.1.P1050125]
Paramecium tetraurelia Cid2 [PTET.51.1.P1340170]
Paramecium octaurelia [POCT.K8.1.P71800002766860159]
Paramecium decaurelia [PDEC.223.1.P00630012]
Paramecium biaurelia [PBIA.V1 4.1.P00410151]
Cid2
Paramecium bursaria Yad1g1n Cid1/3 [TR17851 c0 g1 i11]
Paramecium bursaria CCAP1660/12 Cid1/3 [comp3906 seq2 72 1082]
Paramecium caudatum [PCAU.43c3d.1.P00760077]
Paramecium caudatum [PCAU.43c3d.1.P00410125]
Paramecium jenningsi [PJENN.M.1.P02890016]
Paramecium sexaurelia [PSEX.AZ8 4.1.P1010143]
Paramecium novaurelia [PNOV.TE.1.P06660009]
Paramecium tredecaurelia [PTRED.209.2.P71800001294510023]
Paramecium quadecaurelia [PQUADECEC.NiA.1.P02360007]
Paramecium octaurelia [POCT.K8.1.P71800002769790022]
Paramecium dodecaurelia [PDODEC.274.1.P02140061]
Paramecium tetraurelia Cid3 [PTET.51.1.P0880036]
Paramecium decaurelia [PDEC.223.1.P00990094]
Paramecium biaurelia [PBIA.V1 4.1.P03950017]
Cid1/3
Paramecium sexaurelia [PSEX.AZ8 4.1.P1040135]
Paramecium jenningsi [PJENN.M.1.P02550018]
Paramecium tredecaurelia [PTRED.209.2.P71800001293780009]
Paramecium primaurelia [PPRIMP23072]
Paramecium novaurelia [PNOV.TE.1.P06460013]
Paramecium tetraurelia Cid1 [PTET.51.1.P0910005]
Paramecium octaurelia [POCT.K8.1.P71800002770190015]
Paramecium dodecaurelia [PDODEC.274.1.P03940008]
Paramecium decaurelia [PDEC.223.1.P04000006]
Paramecium biaurelia [PBIA.V1 4.1.P02110014]
Tetrahymena thermophila [EAR92609]
Tetrahymena thermophila [EAR92613]

**Dataset S6.** *Cid* phylogeny (based on 196 aligned sampled amino acid sites) calculated using iqtree with a LG+F+I+G4 best fit substitution model chosen according to Bayesian Inference Criterion (BIC), and with 1,000 non-parametric bootstrap replicates. Classification of *P. bursaria* sequences relative to *Cid1* and *Cid2* is ambiguous, but demonstrates two clear *P. bursaria* orthologues. Three strains of *P. bursaria* (186b, CCAP116/12, & Yad1g1N) were included to improve the phylogenetic resolution of these orthologues. Branches corresponding to *Cid4* and *Cid5* represent two additional Cid orthologues in *Paramecium* that have yet to be described. These orthologues were not detected in any sampled *P. bursaria* or *P. caudatum* datasets, suggesting that they may be exclusive to the *Paramecium aurelia* species complex. Nucleotide sequence (https://doi.org/10.6084/m9.figshare.13387811.v1) and amino acid alignment data (https://doi.org/10.6084/m9.figshare.13387631.v1) for putative *P. bursaria* homologues are available at Figshare.

### Endosymbiont Dcl

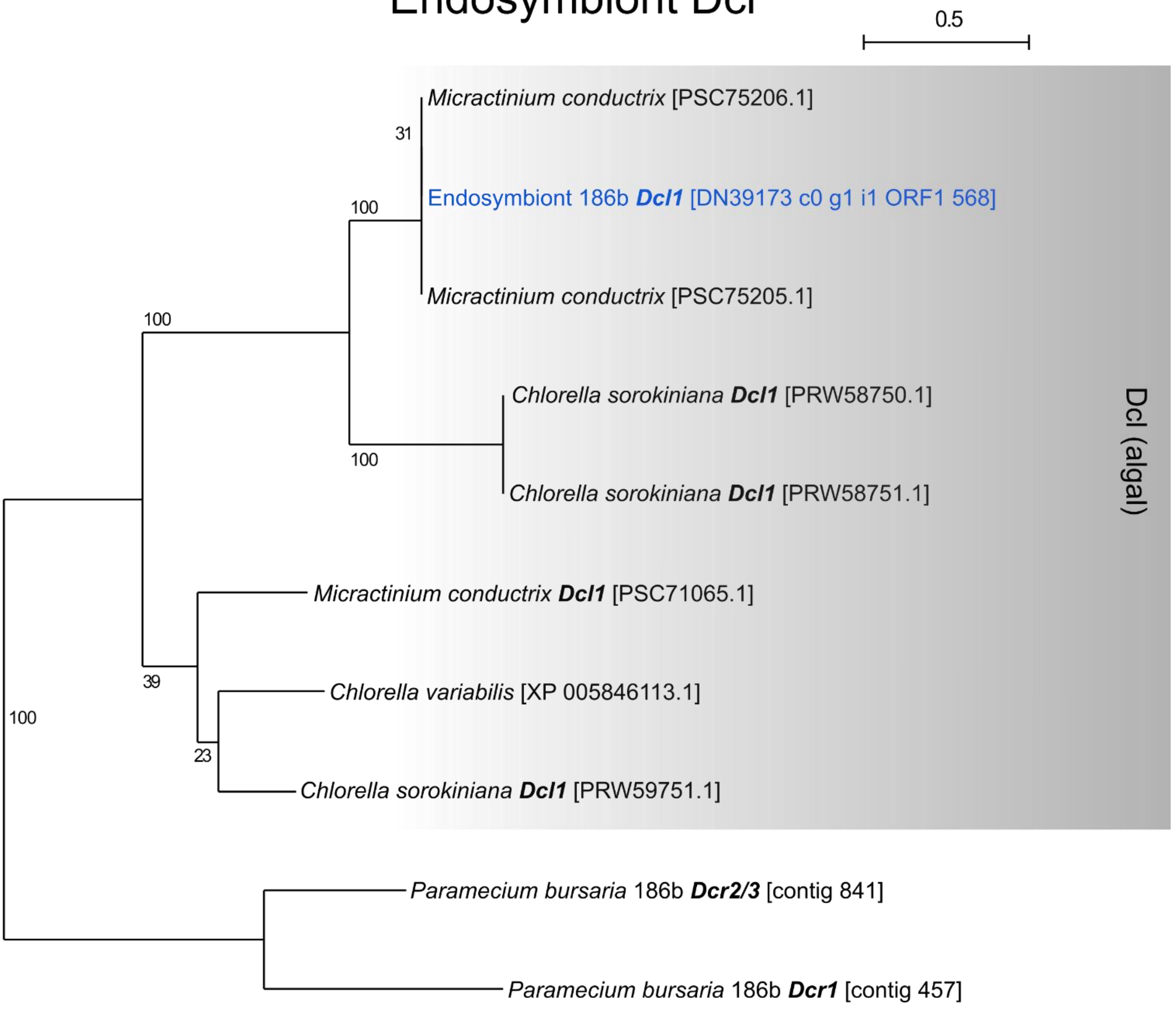


**Dataset S7.** Algal *Dcl* phylogeny (based on 219 aligned amino acid sites) calculated using iqtree with a LG+I+G4 best fit substitution model chosen according to Bayesian Inference Criterion (BIC), and with 1,000 non-parametric bootstrap replicates. Classification of algal Dicer-like sequences demonstrates a single *Dcl1* orthologue in the algal endosymbiont of *P. bursaria* 186b. This sequence is distantly related to the *Dcr1* and *Dcr2/3* orthologues encoded by the *Paramecium* host. Nucleotide sequence (https://doi.org/10.6084/m9.figshare.13387811.v1) and amino acid alignment data (https://doi.org/10.6084/m9.figshare.13387631.v1) for putative *P. bursaria* homologues are available at Figshare.

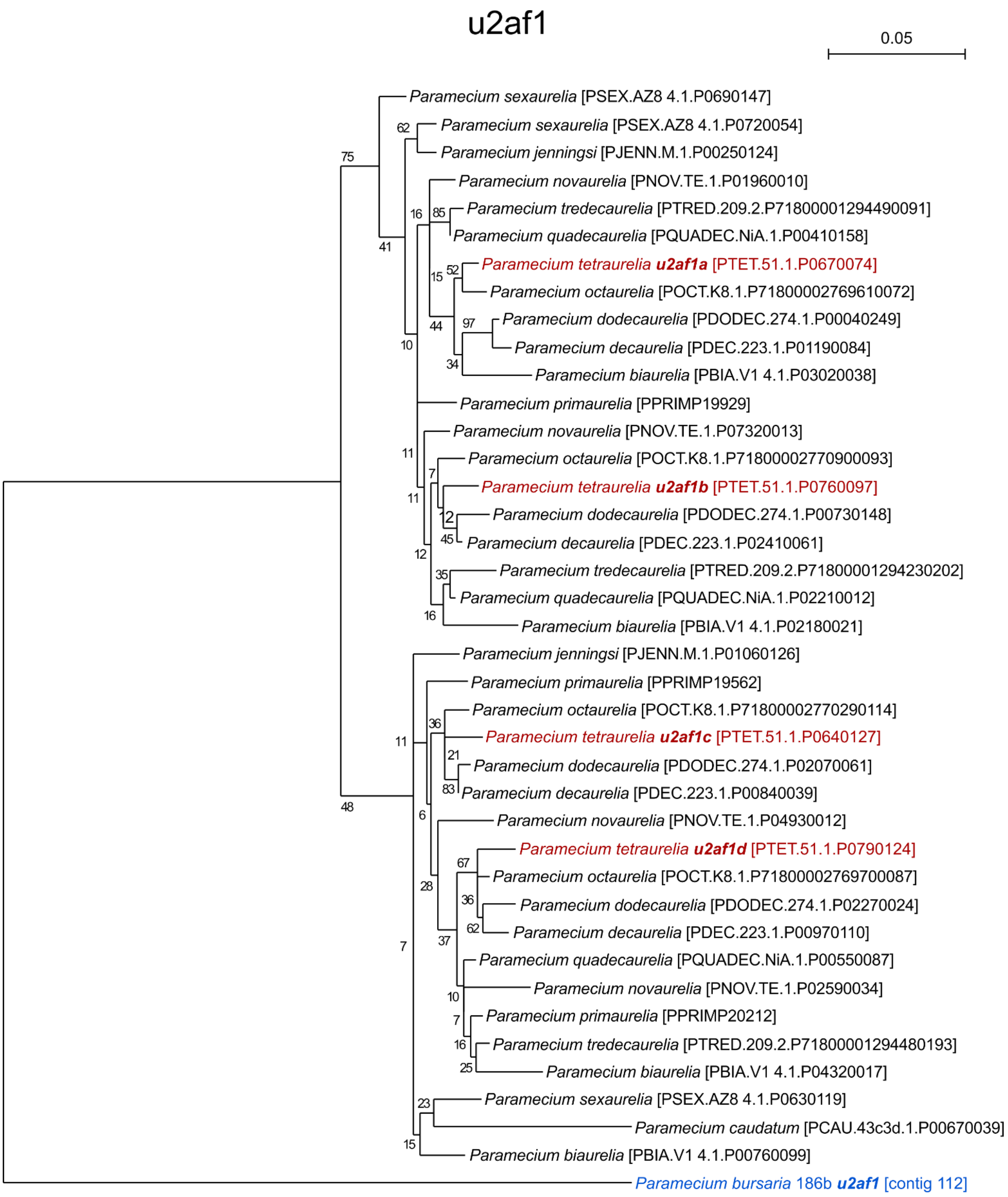


**Dataset S8.** *U2af1* phylogeny (based on 411 aligned amino acid sites) calculated using iqtree with an JTT+R2best fit substitution model chosen according to Bayesian Inference Criterion (BIC), and with 1,000 non-parametric bootstrap replicates. Classification *P. bursaria* sequences demonstrates a single unduplicated *u2af1* orthologue. Note the subsequent duplication of *u2af1* in the *Paramecium aurelia* species complex, with most copies retained. Nucleotide sequence (https://doi.org/10.6084/m9.figshare.13387811.v1) and amino acid alignment data (https://doi.org/10.6084/m9.figshare.13387631.v1) for putative *P. bursaria* homologues are available at Figshare.

#### SUPPLEMENTARY METHODS

##### *Additional transcriptome analysis*

For the purpose of identifying RNAi components in *Paramecium bursaria*, two further *P. bursaria* sequence datasets were analysed. These consisted of a transcriptome sequencing dataset from *P. bursaria* strain CCAP 1660/12 (this study), and the published genome of *P. bursaria* 110224[2]. For the *P. bursaria* CCAP 1660/12 transcriptome dataset, both bulk culture and "single-cell" transcriptomes were generated.

For the bulk transcriptome, ~$10^6$ cell samples were selected 5 hours into the day and 5 hours into the night to represent a 12:12 day-night cycle. RNA extraction was performed using TriReagent (Sigma) following the manufacturer's protocol for extraction from suspension cells, with the addition of an initial lysis step in which samples were heated to 60$^o$C, vortexed with sterile 300 µm glass-beads for 15 s, incubated at room temperature for 10 min, and vortexed for 15s immediately following the addition of TriReagent. Extracted RNA was processed using the Qiagen RNeasy MinElute cleanup kit following the manufacturer's protocol, before being assessed for quality using a NanoDrop ND-1000.

For single cell transcriptomics, a "cell-picking" approach was used in which *P. bursaria* cells (from the CCAP1660/12 culture) were inspected on an inverted light microscope before being picked using an orally aspirated drawn-glass Pasteur[3]. Cell pellets were washed 3 times by serial transfer to 10 µl droplets of sterile NCL media in order to reduce potential bacterial contamination, and transferred to a 10 µL droplet of sterile water. Cells were picked 5 hours into both the light and dark phase of the 12:12 hour day-night cycle identically to the bulk analyses. cDNA was generated and amplified using the MDA-based REPLI-g WTA Single Cell Kit (Qiagen), with additional cell disruption steps consisting of freeze-thaw in liquid nitrogen, bead beating (Sigma, 425-600 µm, acid-washed) and vortexing prior to the addition of lysis buffer. Amplified cDNA was then purified using a QIAamp DNA mini kit (Qiagen). As part of the REPLI-g WTA Single Cell Kit protocol, cDNA was ligated into long fragments by a φ29 DNA polymerase prior to MDA, due to lower MDA efficiency for short fragments[4]. While this approach reduces size-dependent amplification bias, it is important to note that this could potentially lead to the creation of chimeric transcripts in which paired-reads cross boundaries of adjacently ligated cDNA transcripts. For this reason, we conducted additional manual filtering and quality control steps on all *P. bursaria* 'host' and 'endosymbiont' sequence datasets prior to sRNA mapping experiments, even when the respective dataset had not been acquired through this approach. This process of manual filtering is outlined in the **Methods**.

##### *Transcriptome Library Sequencing*

For both bulk and single cell preparations, each cDNA sample was fragmented in 130 µl 1xTE buffer on a Covaris E220 ultrasonicator with a target size of 225 bp (duty factor of 10%, 200 cycles per burst, peak incident power of 175, 200 s at 7°C). Fragment sizes were checked on a BioAnalyzer (Agilent) 7500 DNA chip. cDNA was then concentrated using a GeneRead Size Selection kit (Qiagen) column with an elution in 35 µl. Fragmentation was then repeated 3 times (110 s) until the majority of cDNA in each library was between 200-250 bp.

cDNA ends were then end-repaired, adenylated and adapters ligated using the NEXTFlex (Bioo scientific) sequencing kit according to the manufacturer's instructions. Finally, prepared libraries were size selected using a Blue Pippin machine at a size selection of 350 bp (range 315-385 bp). Individual library concentrations were checked once more on a Bioanalyzer (Agilent).

The bulk day and night libraries were paired-end 76-bp sequenced on separate flowcells using an Illumina Genome Analyzer II. Single cell libraries were multiplexed, and paired-end 150-bp sequenced across two flowcell lanes using an Illumina HiSeq.

***Additional Transcriptome Assembly***

Libraries were screened for contamination by randomly sampling 5 sets of 10,000 PE reads from each library. These samples were then aligned to NCBI's non-redundant 'nr' protein RefSeq database using DIAMOND-BLASTX[5] (at an expectation of 1e-05), and top hits retained. The taxonomic distribution of these hits were recovered using the NCBI taxonomy database. By comparing these distributions between the libraries, and via inspection of GC% distribution, the following libraries were excluded as likely over contaminated:  Dark1-3, Dark1-5, Dark 2-2, Dark 2-7.

All reads were trimmed using Trimmomatic v0.32[6] with adapter clipping (ILLUMINACLIP) and an Q30 minimum sliding window quality-threshold. This threshold was selected as optimal by taking random 5,000 PE reads subsamples from each library, trimming under a broad range of parameter settings, and tallying the number of concordant reads which mapped to a draft assembly.

Libraries were k-mer normalised and low-abundance (likely erroneous) k-mers discarded using the Khmer package[7]. Specifically, reads were interleaved[8] and then digitally normalised using diginorm [9] with a k-mer size and coverage cut-off of 20. Low abundance and likely erroneous k-mers were then filtered relative to the read coverage i.e. low abundance k-mers were removed from high coverage reads but would be more likely to be retained for low coverage reads[10,11].

*De novo* assemblies were generated from the normalised reads using a range of assemblers and parameter settings. Resultant assemblies were compared using standard assembly statistics (e.g. contigs number and size, bases assembled) as implemented in the trinitystats.pl perl script supplied with Trinity[12] and TransRate[13]. Additionally, the reference free probabilistic assembly assessment RSEM-EVAL package (part of DETONATE)[14].

On the basis of these comparisons, the 31-mer Bridger (v2014-12-01)[15] assembly of bulk and taxonomically screened, normalised, k-mer filtered, single cell libraries, was selected as the optimal transcriptome assembly.

***A machine learning approach to Paramecium bursaria CCAP 1660/12 transcriptome binning***

ORFs were called from assembled transcripts using TransDecoder[12] with a minimum protein size of 100 amino acids. *Paramecium* uses an alternative genetic code in which two universal stop codons (UAA, UAG) are reassigned to glutamine. For the purposes of initial binning, ORFs were called and translated using only using the alternative ciliate code. Spuriously extended transcripts were considered favourable to falsely truncated ones. This resulted in 70,095 transcript sequences. A phylogeny-based machine-learning method was developed (https://github.com/fmaguire/dendrogenous) and used to bin the transcripts into 'host', 'endosymbiont', 'food', or 'unknown' bins.

For each transcript ORF sequence, a BLASTP search was performed against a curated 40 genome database. These genomes were selected to cover the sequenced diversity of the tree of life, with a particular focus on green algal and ciliate representatives. Transcripts with more than 5 hits were aligned with Kalign before masking with TrimAL[16]. Any alignment with fewer than 30 sites after masking was then discarded. The masked alignments were used to generate a rapid maximum-likelihood phylogenetic tree with FastTree2[17].

10,000 of these phylogenies were then manually sorted into 'host', 'endosymbiont', 'food', and 'unknown', resulting in a training-set of 2,600 'host', 1,975 'endosymbiont', 3,456 'food', and 1,969 'unknown' transcripts. Phylogenetic features were extracted based on branching distances of the transcript relative to known taxa corresponding to host (e.g. ciliates), endosymbiont (archaeplastida), food (bacteria), and other. This was used to train a K-Neighbours model (via sci-kit learn[18]), which classified the remaining transcripts into the respective bins. From the 70,095 called ORFs in total this process resulted in: 28,050 'host' derived, 673 'endosymbiont', 926 'food' and 40446 'unknown' transcripts.

**REFERENCES**

1. Marker, S., Carradec, Q., Tanty, V., Arnaiz, O. & Meyer, E. A forward genetic screen reveals essential and non-essential RNAi factors in *Paramecium tetraurelia*. *Nucleic Acids Res* **42**, 7268–80 (2014).
2. He, M. *et al.* Genetic basis for the establishment of endosymbiosis in *Paramecium*. *ISME J* **13**, 1360–1369 (2019).
3. Garcia-Cuetos, L., Moestrup, Ø. & Hansen, P. J. Studies on the genus *Mesodinium* II. Ultrastructural and molecular investigations of five marine species help clarifying the taxonomy. *J. Eukaryot. Microbiol.* **59**, 374–400 (2012).
4. Korfhage, C., Fricke, E. & Meier, A. Whole-Transcriptome Amplification of Single Cells for Next-Generation Sequencing. *Curr Protoc Mol Biol* **111**, 7.20.1-7.20.19 (2015).
5. Buchfink, B., Xie, C. & Huson, D. H. Fast and sensitive protein alignment using DIAMOND. *Nat. Methods* **12**, 59–60 (2015).
6. Bolger, A. M., Lohse, M. & Usadel, B. Trimmomatic: a flexible trimmer for Illumina sequence data. *Bioinformatics* **30**, 2114–2120 (2014).
7. Crusoe, M. R. *et al.* The khmer software package: enabling efficient nucleotide sequence analysis. *F1000Res* **4**, 900 (2015).
8. Döring, A., Weese, D., Rausch, T. & Reinert, K. SeqAn An efficient, generic C++ library for sequence analysis. *BMC Bioinformatics* **9**, 11 (2008).
9. Brown, C. T., Howe, A., Zhang, Q., Pyrkosz, A. B. & Brom, T. H. A Reference-Free Algorithm for Computational Normalization of Shotgun Sequencing Data. *arXiv:1203.4802 [q-bio]* (2012).
10. Zhang, Q., Awad, S. & Brown, C. T. Crossing the streams: a framework for streaming analysis of short DNA sequencing reads. https://peerj.com/preprints/890 (2015) doi:10.7287/peerj.preprints.890v1.
11. Zhang, Q., Pell, J., Canino-Koning, R., Howe, A. C. & Brown, C. T. These are not the k-mers you are looking for: efficient online k-mer counting using a probabilistic data structure. *PLoS ONE* **9**, e101271 (2014).
12. Haas, B. J. *et al.* De novo transcript sequence reconstruction from RNA-seq using the Trinity platform for reference generation and analysis. *Nat Protoc* **8**, 1494–1512 (2013).

13. Smith-Unna, R., Boursnell, C., Patro, R., Hibberd, J. & Kelly, S. TransRate: reference free quality assessment of de novo transcriptome assemblies. *Genome Res.* gr.196469.115 (2016) doi:10.1101/gr.196469.115.

14. Li, B. *et al.* Evaluation of de novo transcriptome assemblies from RNA-Seq data. *Genome Biol.* **15**, 553 (2014).

15. Chang, Z. *et al.* Bridger: a new framework for de novo transcriptome assembly using RNA-seq data. *Genome Biol.* **16**, 30 (2015).

16. Capella-Gutiérrez, S., Silla-Martínez, J. M. & Gabaldón, T. trimAl: a tool for automated alignment trimming in large-scale phylogenetic analyses. *Bioinformatics* **25**, 1972–1973 (2009).

17. Price, M. N., Dehal, P. S. & Arkin, A. P. FastTree 2 – Approximately Maximum-Likelihood Trees for Large Alignments. *PLOS ONE* **5**, e9490 (2010).

18. Pedregosa, F. *et al.* Scikit-learn: Machine Learning in Python. *MACHINE LEARNING IN PYTHON* 6.
